# Causal disentanglement for single-cell representations and controllable counterfactual generation

**DOI:** 10.1101/2024.12.11.628077

**Authors:** Yicheng Gao, Kejing Dong, Caihua Shan, Dongsheng Li, Qi Liu

## Abstract

Conducting disentanglement learning on single-cell omics data offers a promising alternative to traditional black-box representation learning by separating the semantic concepts embedded in a biological process. We present CausCell, which incorporates the causal relationships among disentangled concepts within a diffusion model to perform disentanglement learning, with the aim of increasing the explainability, generalizability and controllability of single-cell data, including spatial and temporal omics data, relative to those of the existing black-box representation learning models. Two quantitative evaluation scenarios, i.e., disentanglement and reconstruction, are presented to conduct the first comprehensive single-cell disentanglement learning benchmark, which demonstrates that CausCell outperforms the state-of-the-art methods in both scenarios. Additionally, CausCell can implement controllable generation by intervening with the concepts of single-cell data when given a causal structure. It also has the potential to uncover biological insights by generating counterfactuals from small and noisy single-cell datasets.

## Main

Single-cell technologies have revolutionized the field of biology by enabling analyses to be conducted at the individual cell resolution^1^. This granular perspective has revealed the vast heterogeneity within cell populations, leading to new insights into cellular functions, development processes, and diseases^2^. However, the complexity of single-cell data, which are characterized by high dimensionality and the presence of various entangled concepts, poses great challenges for data interpretation tasks. The process of disentangling these data, i.e., separating complex, intertwined signals into distinct, interpretable components, has therefore emerged as a critical task^3^ and presented to be the necessary path to build the virtual cell^4^.

Disentangled representation learning seeks to capture and separate the underlying concepts embedded in intertwined data, such as images^5^. Unlike traditional end-to-end black-box representation learning methods, which often learn shortcuts by predicting human-annotated labels or reconstructing observable data, disentangled representation learning mimics the human understanding process by leveraging hidden semantic concepts to make decisions^6^. However, this is very challenging for single-cell data, as they are more complex and noisier than data in traditional machine learning communities, such as images. Additionally, these latent concepts in single-cell data are often causally connected, making it hard to clearly separate them with existing disentanglement methods. It requires more advanced techniques to capture latent concepts with causal structure and subsequently establish accurate mappings between the data and concepts in single-cell data.

Several studies have attempted to apply disentangled representation learning to obtain interpretable and manipulable representations of single-cell data. These methods can be categorized into two main groups. (1) Statistical methods: Techniques such as factor analysis or nonnegative matrix factorization are used to identify various biological programs on the basis of statistical patterns^7, 8^. However, these models do not consider the causal structures between concepts and cannot manipulate these concepts. (2) Learning-based methods: These approaches are generative and often utilize variational autoencoder (VAE)-based methods to learn hidden concepts by reconstructing single-cell data^3, 9-11^. For example, CPA decouples perturbation responses^11^, scDisInFact removes batch effects^9^, and Biolord disentangles the concepts contained in single-cell data^3^. However, similar to statistical methods, these methods also cannot guarantee the causality of concepts. In addition, most methods rely on latent optimization^12^, which can result in a loss of fine-grained concept representations at the single-cell resolution level. Notably, prior studies have failed to design and implement comprehensive quantitative benchmarking to rigorously evaluate these disentangled methods. Collectively, a comprehensive benchmarking of existing disentanglement methods, along with the development of a causal disentanglement technique for single cell omics data, is lacking and urgently needed.

Herein, we introduce CausCell, the first deep generative framework for conducting causal disentanglement and counterfactual generation on single-cell omics data (Fig. 1a). CausCell combines a structural causal model^13, 14^ (SCM) with a diffusion model, offering unprecedented advantages in three aspects for obtaining disentangled single-data representations, including for spatial and temporal omics data: (1) Explainability: CausCell leverages an SCM to recover latent concepts with semantic meanings and their causal relationships via a causal directed acyclic graph (cDAG), substantially enhancing the interpretability of the model. (2) Generalizability: Unlike previously developed VAE-based methods, CausCell uses a diffusion model as its backbone, which provides strong generative and generalization capabilities, ensuring a high-quality sample generation process^15^. (3) Controllability: CausCell enables controllable generation by manipulating disentangled representations in the latent space while preserving their consistency with the underlying causal structure. In addition, to disentangle single-cell data into various concepts, we assume that each cell is generated by two types of concepts, i.e., observed and unexplained concepts. For example, observed concepts may involve the cancer type related to single-cell tumour omics data or spatial-domain loci derived from spatial single-cell data, and unexplained concepts are the potential unknown concepts contained in the given data. Therefore, such a framework enables us to effectively distinguish and explore the concepts hidden in the latent space. To train our model, we propose a new loss function that incorporates a new evidence lower bound (ELBO) loss and an independence constraint for the unexplained concepts. As a result, two quantitative evaluation scenarios, i.e., disentanglement and reconstruction, are presented to conduct the first comprehensive single-cell disentanglement learning benchmark, which demonstrates that CausCell outperforms the state-of-the-art tools in both scenarios. Furthermore, we validate the effectiveness of the cDAG in our model, showing that it can generate cells that are consistent with the underlying causal systems when interventions occur. Finally, we show that CausCell can uncover new biological insights when the input experimental single-cell omics dataset is small, noisy and inaccessible. Collectively, CausCell presents an unprecedented perspective and method for obtaining causal disentanglement representations of single-cell data compared with traditional black-box representation learning, and it can uncover novel and interpretable biological insights from single-cell data by controllable counterfactual generation.

**Fig. 1.**
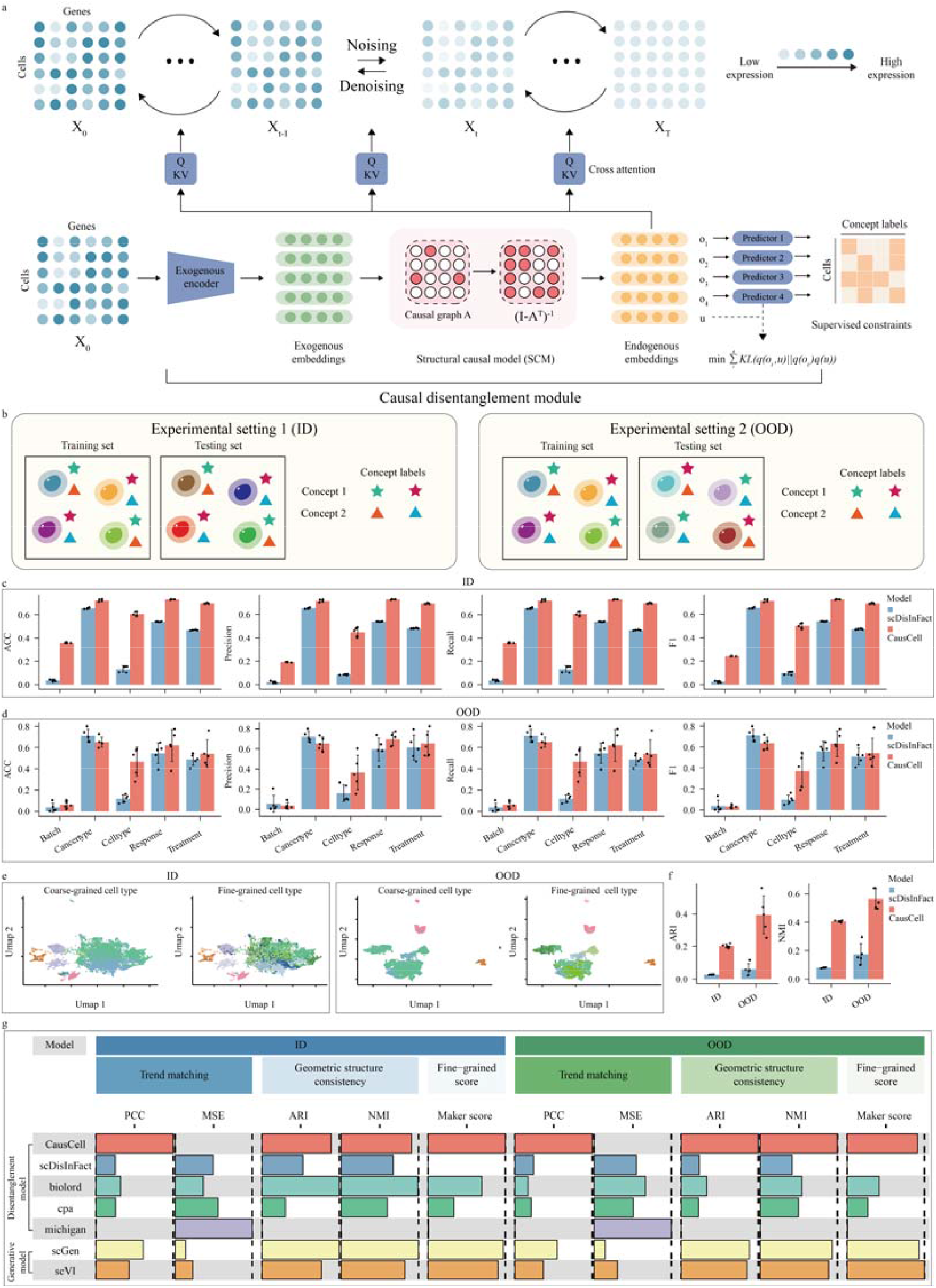
Overview of CausCell and its disentanglement and reconstruction performance. a. The framework of CausCell, which consists of a causal disentanglement module and a diffusion-based generative module. b. The two experimental settings for benchmarking the disentanglement and reconstruction capabilities of the tested model. c. The results of a disentanglement performance comparison conducted under experimental setting 1 (ID) for each concept contained in the ICI_response dataset. d. The results of a disentanglement performance comparison conducted under experimental setting 2 (OOD) for each concept contained in the ICI_response dataset. e. UMAPs of the cell type embeddings produced with coarse-grained and fine-grained cell type annotations across the two experimental settings. f. NMI and ARI metrics attained by different models for evaluating their fine-grained geometric property preservation capabilities across the two experimental settings. g. Comparison among different models in terms of their reconstruction performance across two experimental settings. This is created by the funcky heatmap (version 0.5.0), so the lengths of the bars are proportional to the min-max normalized mean values. For all the bar charts, the data are presented as mean values, with each error bar representing the standard deviation based on n=5 (fivefold cross-validation).

### A comprehensive quantitative disentanglement representation benchmark

A comprehensive quantitative disentanglement model benchmark is critical for establishing the reliability of such models, and this step was overlooked in previous studies. Evaluating the effectiveness of a disentanglement model involves the evaluation of two key aspects: (1) concept disentanglement and (2) reconstruction. The first aspect reflects the ability of the tested model to accurately capture and separate underlying semantic concepts, whereas the second aspect determines the quality and fidelity of the generated counterfactual samples by manipulating concepts for further biological analysis. Properly evaluating these aspects is essential for ensuring the practical applicability and robustness of the developed model.

To conduct comprehensive benchmarking, we collected five distinct single-cell datasets spanning different biological domains^16-20^, each with various causal relationships among different concepts (Supplementary Note 1 and 2, Supplementary Fig. 1 and Supplementary Table 1). We also included two experimental settings, in-distribution (ID) and out-of-distribution (OOD) settings, on the basis of whether the different combinations of concept labels were presented during training (Fig. 2b and Supplementary Note 3). In the ID setting, the model had seen all possible combinations of concept labels during training, providing a direct assessment of its ability to operate within the same data distribution. The more challenging OOD setting involved cases where the model encountered unseen combinations of concept labels, and this task aimed to assess the ability of the model to transfer complex relationships among concept representations and examine its generalizability. Furthermore, we defined five types of metrics to evaluate various aspects of disentanglement and reconstruction performance (Supplementary Note 4). Disentanglement performance was evaluated by determining whether the learned concept representation contained sufficient information for predicting the concept labels. Therefore, we defined (1) the predictive ability of the concept embeddings and (2) the clustering consistency of the embeddings when the label granularity was varied (Supplementary Note 5). For reconstruction, we defined (3) trend matching, (4) geometric structure consistency and (5) a fine-grained score to test the consistency of the generated single-cell data with the test data distribution.

**Fig. 2.**
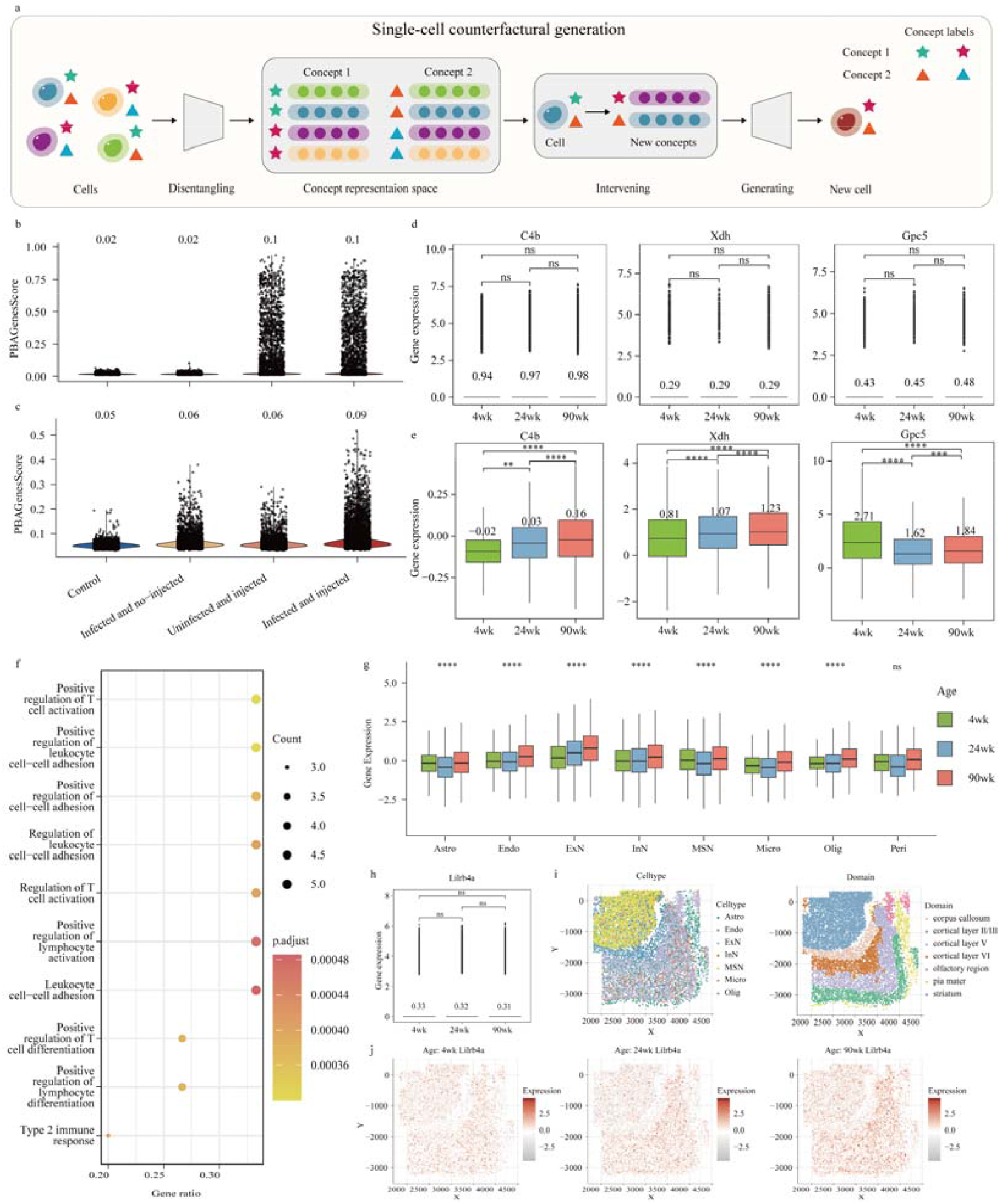
The counterfactual generation results produced by CausCell. a. Schematic diagram of the single-cell counterfactual generation process implemented by CausCell. b. PBAGenesScore values obtained for various intervention-associated cells generated by CausCell without incorporating a causal structure (CausCell_IND). c. PBAGenesScore values produced for various intervention-associated cells generated by the CausCell version incorporating the causal structure. d. Gene expression differences among different ages in the original dataset, where *C4b* in Oligo, *Xdh* in Endo and *Gpc5* in Astro. e. Gene expression differences among different ages in the counterfactually generated dataset, where *C4b* in Oligo, *Xdh* in Endo and *Gpc5* in Astro. f. GO enrichment analysis of the differentially expressed upregulated genes in microglia enriched in the striatum domain. g. Age-related *Lilrb4a* gene expression differences across all cell types. h. *Lilrb4a* expression differences among different ages in the original dataset. i. Spatial segmentation results concerning the anatomical regions of the mouse brain. j. The spatial distribution characteristics of *Lilrb4a* expression across all ages. In the GO enrichment analysis, a hypergeometric test was performed. In each boxplot, the box boundaries represent the interquartile range, the whiskers extend to the most extreme data points within 1.5 times the interquartile range, the value indicated above the box represents the mean value, and the black line inside the box represents the median. Statistical tests were conducted via a two-sided t test with significance levels of ns: p>0.05; ^*^: p≤0.05; ^**^: p≤0.01; ^***^: p≤0.001; ^****^: p≤0.0001), except for subfigure G, where an analysis of variance (ANOVA) test was used.

To evaluate the disentanglement performance of the proposed approach, scDisInFact was selected as the baseline model because it uniquely maintains a single-cell resolution in concept representation tasks. Most existing disentanglement models rely on latent optimization, wherein cells sharing the same concept label are represented by identical concept representations^12^. These approaches compromise the single-cell resolution, thereby hindering the disentanglement performance assessment. Preserving concept representations at the single-cell level is crucial, as doing so captures the heterogeneity inherent within cell populations. For example, within the T-cell lineage, distinct subtypes, such as exhausted T cells and effector T cells, exhibit unique functional states. Utilizing a uniform representation for all T cells would obscure these subtle biological distinctions. Our results showed that CausCell outperformed scDisInFact in terms of various predictive metrics, including accuracy, precision, weighted F1 scores and weighted recall scores (Figs. 1c, d and Supplementary Figs. 2-5). To evaluate the concept embedding generalization capabilities of the models, we varied the granularity of their concept labels by training the models on coarse-grained cell type information and subsequently assessing their performance in terms of fine-grained cell subtype consistency (Supplementary Note 6). CausCell also demonstrated superior performance across several clustering consistency metrics, such as the normalized mutual information (NMI) and adjusted Rand index (ARI) scores (Figs. 1e, f and Supplementary Note 7).

To evaluate the reconstruction performance of the models, we used the Pearson correlation coefficient (PCC) and mean squared error (MSE) to measure the consistency of the trend between the generated data and the original data. Additionally, NMI and the ARI were employed to assess the preservation of geometric structures. A fine-grained score was also introduced to evaluate the maintenance of the marker genes in the generated cells. Then, CausCell was benchmarked against six baseline models, including four mainstream disentanglement-based models (Biolord^3^, scDisInFact^9^, CPA^11^ and MichiGAN^10^) and two generative models without disentanglement (scVI^21^ and scGen^22^) across all the above evaluation metrics. Theoretically, a trade-off exists between disentanglement and reconstruction performance^23, 24^. Our results demonstrated that CausCell not only surpassed all disentanglement-based models but also achieved reconstruction performance that was on par with or superior to that of the mainstream generative models (Fig. 1g and Supplementary Figs. 2-5). The exceptional performance of CausCell in both disentanglement and reconstruction tasks provided a solid foundation for various downstream analyses. These included generating samples that adhered to causal structures and producing reliable counterfactual samples, thereby facilitating the discovery of new biological insights.

### Realistic biological application of CausCell

First, we demonstrated that CausCell could enhance the plausibility and consistency of the counterfactual generation process by aligning with the underlying causal structure (Fig. 2a). Counterfactual generation plays a critical role in generating hypothetical versions of single-cell data by intervening in one or several concepts, and it aims to investigate how scientific conclusions change and explore the reasons behind these changes, presenting to be an important task to build a virtual cell^4^. However, the existing disentanglement models exhibit a critical limitation because they intervene in concepts without considering their causal relationships with other concepts, resulting in the generation of unrealistic or erroneous samples. To further illustrate this point, we compared the quality levels of the samples generated by CausCell and a modified version, CausCell_IND, which lacks the causal structure and treats concept relationships as independent by removing the SCM layer from the model. Utilizing the “Control” cells contained in a spatial-temporal single cell data set, i.e., the Spatiotemporally_liver dataset, we performed interventions on the “Inject,” “Time,” and “Infect” concepts by modifying their concept labels and generating counterfactual cells with modified concept labels. We then compared the infection statuses of cells quantified by their PBAGenesScore^18^ values (Supplementary Note 8). Regarding CausCell_IND, we observed that the cells generated with the “Inject” intervention presented the highest PBAGenesScore values across all time points, regardless of whether the cells were infected, even when the time value was set to 0 (Fig. 2c and Supplementary Fig. 6). This unrealistic behaviour was mitigated by incorporating the causal structure into CausCell, allowing the model to generate more realistic samples that were consistent with the underlying causal relationships, where the injected and infected cells presented higher scores than other conditions did (Fig. 2b and Supplementary Fig. 6).

Second, we validated that CausCell could reveal biological insights even from small and noisy single-cell datasets via controllable counterfactual generation (Supplementary Note 9). Single-cell data are often confounded by latent concepts, which may obscure critical biological signals, especially when the sample size is small. As a result, wet-lab experiments typically require large sample sizes and significant costs for high-throughput sequencing. The controllable counterfactual generation process conducted by CausCell offers a promising solution for augmenting data when the number of data samples is small. To illustrate this point, we analysed another spatial-temporal dataset, i.e., the mouse ageing dataset (MERFISH_brain)^17^, which includes 3 mice aged 4 weeks (4wk), 3 mice aged 24 weeks (24wk), and 5 mice aged 90 weeks (90wk). A previous study^17^ revealed that the expressions of 3 genes in oligodendrocytes (Oligo), 2 genes in microglia (Micro), 2 genes in astrocytes (Astro) and 1 gene in endothelial cells (Endo) increased with age, whereas the expression of one gene (*Gpc5*) in Astro decreased. However, when the sample sizes were reduced (by selecting only 2 mice from each age group), these gene expression trends were no longer observed (Fig. 2d and Supplementary Fig. 7). We subsequently used this smaller dataset to train CausCell and performed controllable counterfactual generation on the 4-week-old mice by intervening in the age concept to simulate mice at 24 and 90 weeks of age. By using these counterfactual samples, we replicated the analysis and found that 6 out of the 9 genes presented the same expression trends as previous study^17^ (Fig. 2e and Supplementary Fig. 7). Additionally, the abundance of cells with high degrees of expression for 2 of the remaining 3 genes also increased (Supplementary Fig. 8).

Furthermore, we statistically analysed the number of age-related differential genes contained in different cell types across various spatial domains, which yielded results similar to those of a previous study^17^ (Supplementary Fig. 9 and Supplementary Note 9). Notably, we obtained a new finding: the upregulated genes in Micro were enriched in the striatum domain, which has never been reported in previous studies^17^. A GO enrichment analysis^25^ revealed that these age-related, upregulated genes were associated with multiple cell adhesion pathways, particularly those involving immune cells, in addition to immune activation pathways (Fig. 2f and Supplementary Note 10). These findings suggest that Micro may contribute to brain ageing through cell adhesion mechanisms. A further analysis identified *Lilrb4a* as the gene with the most enriched pathways, including T-cell activation and leukocyte adhesion, which were not highlighted in earlier work and could not be identified in the original data (Fig. 2h). A recent study^26^ revealed that Micro in Alzheimer’s disease patients express high levels of *LILRB4*, the homologous gene to *Lilrb4a*. Additionally, its family gene *LilrB3* can activate microglia into a proinflammatory state^27^. This finding highlights the potential role of *Lilrb4a* in influencing brain ageing by regulating the immune states of microglia cells. Finally, we assessed the expression of *Lilrb4a* across different cell types and spatial domains and found that its expression increased with age in both neuronal and nonneuronal cells, with spatial specificity (Figs. 2i, j). These findings suggest that *Lilrb4a* may be a common regulator of ageing across various cell types in the brains of mice.

## Discussion

In conclusion, we presented comprehensive evaluation metrics that demonstrate the superior disentanglement and reconstruction performance of CausCell in single-cell disentanglement representation tasks. Additionally, by leveraging causal structures and diffusion models, CausCell has the potential to generate more realistic samples and uncover meaningful biological insights through the generation of counterfactuals, particularly when the given sample size is small. Future updating of CausCell are expected: (1) Herein, we applied the linear version of an SCM, i.e., we utilized a closed-form solution to transform concept representations given a predefined causal structure. In practice, the relationships among concepts can be nonlinearly defined in a causal structure. We need to add more learnable SCM layers to transform concepts and represent their causal relationships. (2) While our focus was on evaluating the performance of the disentanglement model and demonstrating the benefits of the causal prior in the counterfactual generation task, many potential applications of this model remain to be explored. The current CausCell framework can be naturally extended to model more modalities in single-cell data, even a foundational disentanglement model in the future.

## Methods

### Overview of CausCell

Consider the input data a single-cell sequencing dataset of *N* cells 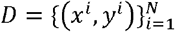, where *x*^*i*^ represents the gene expression vector for cell *i* and *y*^*i*^consists of *M* observed concept labels for that cell. The disentanglement process involves learning a series of latent concept embeddings 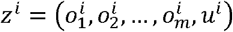, where each 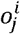 is a low-dimensional vector corresponding to the label 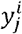, and *u*^*i*^ serves as an extra captured embedding of the unobserved concepts. These embeddings *z*^*i*^ are then employed during the generative process to achieve a better reconstruction effect. Previously, VAEs were implemented as generative modules, but they tend to produce lower-quality samples, especially when addressing high-dimensional data such as high-resolution images data^28^. In contrast, diffusion models have been demonstrated to produce high-quality samples by generating samples in a step-by-step manner. However, their lack of an interpretable latent space makes them less suitable for disentanglement and explainability purposes. To address these issues, we propose CausCell to learn a causal disentanglement module *F* for concept embeddings and then integrate them into the diffusion process *G*. Both modules are incorporated into a unified network, which is end-to-end trained via a newly derived ELBO loss.

### Causal disentanglement module *F*

The goal is to learn causal representations of concepts *z*^*i*^. The disentanglement learning methods used in previous single-cell studies were constrained by the assumption of concept independence. However, concepts are not necessarily independent in practice. Instead, an underlying causal structure is present and renders these concepts dependent. Therefore, we introduce an SCM layer to construct causal relationships among the concepts. Specifically, the observed concepts 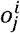 are connected through a DAG, which is represented by an adjacency matrix A. We apply a linear version of the SCM, which satisfies the following equation:

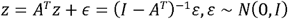

where *ε* represents the exogenous variables and *z* denotes the endogenous variables for capturing the latent concepts. We first learn a function (a multilayer perceptron (MLP)) for extracting the exogenous concept embeddings *ε* from the gene expression *x*_0_ and then leverage *ε* to transform it into *z* via a closed-form solution.

To ensure the semantic meanings of the concept embeddings and to enforce causal disentanglement, we employ *m* discriminators *D* = {*D*_1_, *D*_2_,…, *D*_*m*_}, each of which corresponds to an observed concept. These discriminators are trained to predict the observed concept labels from the embeddings 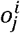 via the cross-entropy loss:

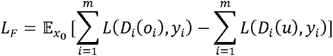

where *L* is the cross-entropy loss function. This setup also encourages independence between the observed concepts *o* and the unexplained concept *u*.

### Generative module *G*

We use a diffusion model as our generative backbone in CausCell because of its powerful generative capabilities. A diffusion model defines a latent variable distribution *p*(*x*_0:*T*_) over a gene expression vector x_0_ sampled from a single-cell distribution, as well as noisy gene expression vectors *x*_1:*T*_ ≔ *x*_1_, *x*_2_,…, *x*_*T*_ that represent a gradual transformation of *x*_0_ into random Gaussian noise *x*_*T*_. The reverse diffusion process is modelled as a Markov chain:

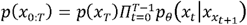

where *p*_*θ*_ is a learned denoising distribution parameterized by a neural network with a parameter *θ*.

The forward diffusion process *q* adds Gaussian noise to *x*_0_ at each step:

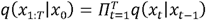

with a predefined noise schedule 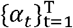. The marginal distribution can be directly computed as shown below:

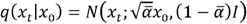

where 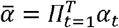.

The denoising network *g*_*θ*_ in the diffusion model is an *ϵ* predictor in most cases. However, single-cell data often exhibit extreme sparsity, and the corrupted input at time step *t* is mostly pure noise. Under this setting, the model is likely to learn to reverse the noise schedule instead of the true data posterior. Therefore, we adopt the *x*_0_-predictor^29^ in this study, and a simplified training loss for the generative module is defined as follows:

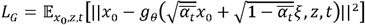

where *ξ* is the noise sampled from *N* ∼ (0, *I*) and *t* is the time step.

### Integration causal disentanglement module *F* with a diffusion process *G*

The causal disentanglement module takes a single-cell expression profile as its input to obtain a set of concept embeddings *z*^*i*^ under the guarantee of a causal structure. We then use these concept embeddings as the condition information for the generative module. Specifically, we incorporate these concept embeddings into the reverse diffusion process of the generative model through a cross-attention mechanism. By conditioning this process on the concept embeddings, the model can generate gene expression profiles that are consistent with specific biological concepts, allowing for a controlled and interpretable data generation procedure. The cross-attention mechanism^30^ facilitates this conditioning task by enabling the model to focus on the relevant parts of the concept embeddings during the generation phase. It is mathematically defined as:

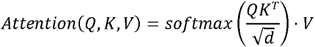

where *Q, K*, and *V* are the query, key, and value vectors, respectively. *d* is the scaling vector, which is used to ensure numerical stability in the softmax function, and its value is set as the dimensionality of the head. In this study, the corrupted gene expression profile 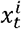 in the diffusion module serves as a query, and the concept embeddings *z*^*i*^ act as keys and values.

### Evidence lower bound of CausCell

We formulate the training objective via a variational inference approach to derive the ELBO for optimizing the model parameters. We treat both *z* and *ϵ* as latent variables. Consider the following conditional generative model:

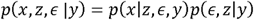

We define *p*_*ϵ*_(*ϵ*) = *N*(0,*I*) and the joint prior *p*(*ϵ,z*|*y*) for latent variables *z* and *ϵ* as follows^14^:

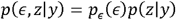

We define CausCell as a causal disentanglement-based diffusion model represented by the conditional probability distribution *p* (*x*_0:*T*_, *z, ϵ*| *y*), which can be factorized as follows:

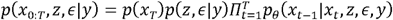

This model implements a reverse diffusion process *p*_*θ*_(*x*_*t*−1_|*x*_*t*_,*z,ϵ,y*) over *x*_0:*T*_, which is conditioned on the endogenous variables *z*, the exogenous variables *ϵ* and the concept labels y. All of these variables are independent of the diffusion process because these variables are properties of the input, not control variables of the diffusion process.

To optimize the model parameters, we apply variational inference twice to impose a variational lower bound on the conditional log-likelihood of the concept labels *log p*(*x*_*o*_|*y*):

#### Proposition 1

The ELBO of CausCell can be derived as follows (Proof: Supplementary Note 11):

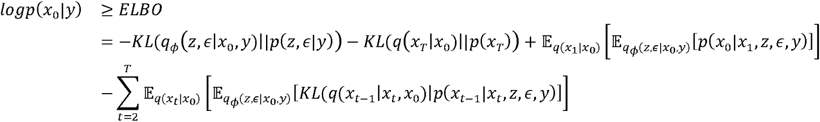

The above equation is intractable in general. Given an SCM with latent exogenous independent variables *ϵ* and the latent endogenous variables *z*, we have *z* = *A*^*T*^*z* + *z ϵ* = (*I* − *A*^*T*^)^−1^*ϵ*. For simplicity, we denote *C* = (*I* − *A*^*T*^)^−1^. Leveraging the one-to-one correspondence between *ϵ* and *z*, we can simplify the variational posterior as follows:

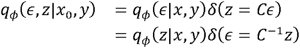

where *δ*(·) is the Dirac delta function. According to the model assumptions introduced above, we can further simplify the ELBO as follows:

#### Proposition 2

The ELBO of CausCell can be rewritten as follows (Proof: Supplementary Note 12):

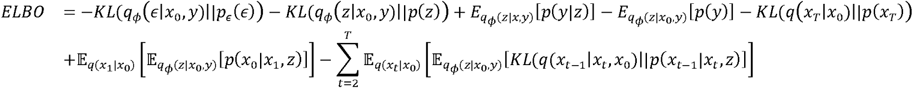

In practical scenarios, the latent factors *z* consist of observed variables *o* and unobserved variables u such that *z* = [*o,u*]. Assuming independence between *o* and *u*, we augment the ELBO with a term 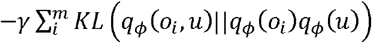 that encourages this independence, and it is implemented via an adversarial debiasing strategy^31^ using discriminators. Therefore, the overall training objective is to maximize the ELBO, leading to the following combined loss function (Supplementary Note 14):

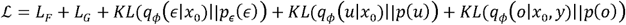

### Benefit of disentanglement in the diffusion model

We show that the reverse diffusion process introduces an information bottleneck effect, promoting disentanglement by dynamically allocating information to the latent concepts as the time steps increases^32^. This is reflected in the ELBO term for the reverse diffusion process, which can be formulated as follows.

#### Proposition 3

The reverse diffusion process term in the ELBO can be rewritten as shown below (Proof: Supplementary Note 13):

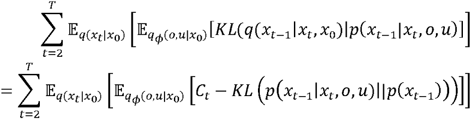

where *C*_*t*_ denotes the Kullback-Leibler (KL) divergence between the determined distribution *q*(*x*_*t*−1_|*x*_*t*_,*x*_0_) and the standard Gaussian distribution *p*(*x*_*t*−1_) := *N*(0, *I*).

Therefore, optimizing the model encourages the KL divergence *KL* (*p*(*x*_*t*−1_|*x*_*t*_,*o,u*)||*p*(*x*_*t*−1_)) to approximate a constant *C*_*t*_, effectively regulating the information content of *x*_*t*−1_. The larger the KL divergence is, the more information *x*_*t*−1,_ carries. By promoting *KL* (*p*(*x*_*t*−1_|*x*_*t*_,*o,u*)||*p*(*x*_*t*−1_)) to approximate *C*_*t*_, an information bottleneck effect is added to it and thus transferred to the latent factors *o, u*. Therefore, the diffusion model has a natural information bottleneck and is a good inductive bias for disentanglement representation purposes^32^. As different concepts to be disentangled may contain different amounts of information, the diffusion model can dynamically allocate this information to the latent concepts in the reverse diffusion step, where the information decreases as the number of time steps increases. This optimization objective is then similar to that of AnnealVAE^33^.

### Single-cell counterfactual generation via the do operator

CausCell performs interventions on individual cells by modifying their associated concepts, enabling the generation of counterfactual gene expression profiles. The causal disentanglement module of CausCell maps the concept representations of all *N* cells to a concept representation space *Ω*. In this space, each concept *o*_*i*_ has a representation for every cell, resulting in a total of *N* representations per concept. For each concept *o*_*i*_, there exist *m* distinct concept labels labelled *a*_1_,*a*_2_, …, *a*_*m*_, each of which is associated with a subset of cells such that the total number of cells across all concept labels equals *N*.

To perform a concept intervention on a specific cell *c* that originally had a concept label of *a*_1_ for concept *o*_*i*_, we apply the do operator to set the concept to a new value *a*_*k*_, which is denoted as *do*(*o*_*i*_ = *a*_*k*_). This intervention is implemented in three steps. (1) Randomly select a concept representation from the set of representations associated with the concept label *a*_*k*_ for concept *o*_*i*_. (2) Replace the original concept representation of cell *c* with the selected representation. (3) Keep the representations of all other concepts *o*_*j*≠*i*_, including any unexplained concepts, unchanged for cell *c*. The postintervention distribution is formally represented as follows:

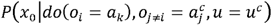

where 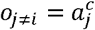denotes that all other concepts *o*_*j*_ (for *j* ≠ *i*) retain their original concept labels 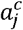, as in the original cell *c*. Utilizing these intervened concept representations, the generative diffusion module of CausCell generates a counterfactual gene expression profile *ĉ* by sampling from the postintervention distribution:

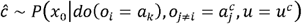

This process allows us to simulate how the gene expression profile of cell *c* would appear under an intervention *do*(*o*_*i*_ = *a*_*k*_), isolating the causal effect of changing concept *o*_*i*_ from *a*_1_ to *a*_*k*_ while controlling for all other concepts.

## Data availability

All datasets utilized by CausCell for training, testing, and application examples were obtained from publicly accessible databases. Specifically, the Immun_atlas dataset was sourced from (https://www.tissueimmunecellatlas.org), the MERFISH_Brain dataset from (https://cellxgene.cziscience.com/collections/31937775-0602-4e52-a799-b6acdd2bac2e), and the Spatiotemporally_Liver dataset from (https://zenodo.org/records/7081863). The ICI_Response dataset was retrieved from the Gene Expression Omnibus (GEO) with accession number GSE123814, while the Limb_development dataset was accessed from ArrayExpress with accession ID E-MTAB-10514.

## Acknowledgements

This work was supported by National Natural Science Foundation of China (Grant No. T24250193, 32341008), the National Key Research and Development Program of China (Grant No. 2021YFF1201200, No. 2021YFF1200900), Shanghai Pilot Program for Basic Research, Shanghai Science and Technology Innovation Action Plan-Key Specialization in Computational Biology, Shanghai Shuguang Scholars Project, Shanghai Excellent Academic Leader Project, Shanghai Municipal Science and Technology Major Project (Grant No. 2021SHZDZX0100) and Fundamental Research Funds for the Central Universities. This work was partially funded by Microsoft Research Asia.

## Author Contributions Statement

Qi Liu, Yicheng Gao and Kejing Dong designed the framework of this work. Dongsheng Li and Caihua Shan provided technical support. Yicheng Gao, Kejing Dong performed the analyses. Yicheng Gao, Kejing Dong, Caihua Shan and Qi Liu wrote the manuscript with the help of other authors. All authors read and approved the final manuscript.

